# Single-cell RNA-seq reveals that glioblastoma recapitulates normal brain development

**DOI:** 10.1101/449439

**Authors:** Charles P. Couturier, Shamini Ayyadhury, Phuong U. Le, Jean Monlong, Gabriele Riva, Redouane Allache, Salma Baig, Xiaohua Yan, Mathieu Bourgey, Changseok Lee, Yu Chang David Wang, V. Wee Yong, Marie-Christine Guiot, Bratislav Misic, Jack Antel, Guillaume Bourque, Jiannis Ragoussis, Kevin Petrecca

**Affiliations:** Department of Neurology and Neurosurgery, Montreal Neurological Institute and Hospital, McGill University, Montreal, QC, Canada; Department of Human Genetics, McGill University, Montreal, QC, Canada; Canadian Centre for Computational Genomics, McGill University, Montreal, QC, Canada; McGill University and Genome Québec Innovation Centre, Montreal, Qc, Canada; Department of Clinical Neurosciences, University of Calgary, Calgary, AB, Canada; Department of Neuropathology, Montreal Neurological Institute and Hospital, McGill University, Montreal, QC, Canada; Department of Bioengineering, McGill University, Montreal, QC, Canada; Lead Contact

**Keywords:** cancer stem cells, glioblastoma, single-cell RNA-seq, heterogeneity, hierarchy, roadmap, brain development

## Abstract

**Summary:** Cancer stem cells are critical for cancer initiation, development, and resistance to treatments. Our understanding of these processes, and how they relate to glioblastoma heterogeneity, is limited. To overcome these limitations, we performed single-cell RNA-sequencing on 38 296 glioblastoma cells and 22 637 normal human fetal brain cells. Using an unbiased approach, we mapped the lineage hierarchy of the developing human brain and compared the transcriptome of each cancer cell to this roadmap. We discovered a conserved neural trilineage cancer hierarchy with glial progenitor-like cells at the apex. We also found that this progenitor population contains the majority of cancer’s cycling cells and is the origin of heterogeneity. Finally, we show that this hierarchal map can be used to identify therapeutic targets specific to progenitor cancer stem cells. Our analyses show that normal brain development reconciles glioblastoma development, unravels the origin of glioblastoma heterogeneity, and helps to identify cancer stem cell-specific targets.

## Introduction

Significant obstacles hampering the development of effective cancer therapeutics include tumour heterogeneity ^1–5^, and the persistence of poorly understood cancer stem cells (CSCs) that give rise to cancer recurrence ^6, 7^.

Glioblastoma, the most common adult brain cancer ^8^, exemplifies these obstacles. Following radiotherapy and temozolomide chemotherapy, the median time to recurrence is 7 months, with patients succumbing to the disease 7 months thereafter ^9, 10^. This cancer is composed of 2 main cell compartments: a larger differentiated cell compartment that forms the basis of our understanding of the genomic and molecular underpinnings of the disease ^11, 12^; and a smaller, less well characterized compartment of cells with stem-like capabilities ^13–16^. The molecular and genomic heterogeneity within the differentiated cell compartment, and the persistence of a subpopulation of cancer cells with stem-like properties following radiotherapy and chemotherapy, are the main causes of resistance to treatment and the associated extremely poor outcomes ^6, 17, 18^.

Interpatient heterogeneity was established through genomic and transcriptomic analyses by The Cancer Genome Atlas (TCGA) research network ^11^. Analysis of whole tumour transcriptomic data extracted from predominantly differentiated cells showed that glioblastoma clustered into 4 main subtypes: proneural; neural; classical; and mesenchymal ^19^. Despite very different transcriptomic profiles and associated genomic alterations, no differences in survival exist between these subtypes. More recently, it has been shown that multiple subtypes coexist in different regions ^20^ and different cells ^12^ within the same tumour. This interpatient and intratumoural heterogeneity poses a daunting challenge for research programs aimed at developing targeted therapeutic approaches ^21^ and may explain the failures of such approaches in this disease.

Another layer of complexity was uncovered by the discovery of a small subpopulation of glioblastoma cells that have stem-like properties ^13, 14^. The cancer stem cell theory is derived from our understanding of normal stem cells ^15^ and posits that such cells must exhibit properties of self-renewal and the ability to produce differentiated progeny. Consistently, glioblastoma stem cells (GSCs) do possess these properties. GSCs can propagate tumours from one host to another ^14, 22^, and can expand and develop to form brain cancers in orthotopic xenograft models that recapitulate the tumour from which they were extracted ^14, 23^. Importantly, stem cells isolated from different tumours show variability with respect to marker expression ^18, 24, 25^, suggesting that some degree of interpatient and/or intratumoural heterogeneity exists within the stem cell compartment as well. While the GSC compartment is small in comparison to the differentiated compartment, it is relevant clinically. Studies have shown that GSCs resist radiotherapy ^6^ and temozolomide chemotherapy ^18, 26^. These data suggest that GSCs may play a role in cancer development and recurrence. There are presently no treatments targeting GSCs.

Our understanding of glioblastoma heterogeneity development, and the relevance of GSCs in this process, is limited. Here, using massively parallel single-cell RNA sequencing (scRNAseq) of glioblastoma and the normal developing human brain, we discovered a conserved trilineage cancer hierarchy with progenitor cancer cells at the apex of this hierarchy. We found that this progenitor population contains the majority of the cancer’s cycling cells, is the origin of glioblastoma heterogeneity, and functionally corresponds to GSCs. Therapeutically relevant, we show that this hierarchal map can be used to identify therapeutic targets specific to GSCs.

## Results

### Single-cell RNA sequencing highlights genomic and transcriptomic heterogeneity in glioblastoma

We used droplet-based scRNAseq ^27–29^ to obtain the transcriptome of cells isolated from freshly-excised glioblastoma and freshly-derived enriched GSCs. Cells were dissociated from the whole tumour and cDNA libraries were prepared on the operative day (Supplementary Fig. 1a). Enriched GSCs were obtained by culturing cancer cells in restricted media ^16^ for one week followed by cDNA library preparation (Supplementary Fig. 1a). GSC lines were proven to be tumourigenic by xenotransplantation. In total, 38 296 cells from 8 patients (Supplementary Fig. 1b) were sequenced: 18 334 whole tumour cells and 20 062 enriched-GSCs.

To distinguish cancer cells from normal brain cells we determined the main copy number aberration (CNA) events in each cell from its transcriptomic profile. Briefly, for every cell in our datasets, we averaged the expression values of adjacent genes on the genome, as has been previously reported ^30–32^. The expression of neighbouring genes should be independent; however, shared CNAs will cause them to vary together on average. CNAs are thus detectable when averaging over a sufficiently large genomic region. We applied the Louvain algorithm on this location-averaged expression data to cluster cells based on common CNAs (Fig. 1a and Supplementary Fig. 1c). Two small clusters devoid of known recurrent CNAs, and containing cells from multiple tumours, were identified (Fig. 1b and Supplementary Fig 1d). Cells in these clusters expressed genes found exclusively in lymphocytes, oligodendrocytes, or endothelial cells (Fig. 1c), and were thus classified as normal cells. All other clusters were formed by cells originating from a single tumour and contained multiple CNAs. We defined these as cancer cells. When enriched GSCs and whole tumour cells were sequenced from the same patient these samples clustered together.

**Figure 1.**
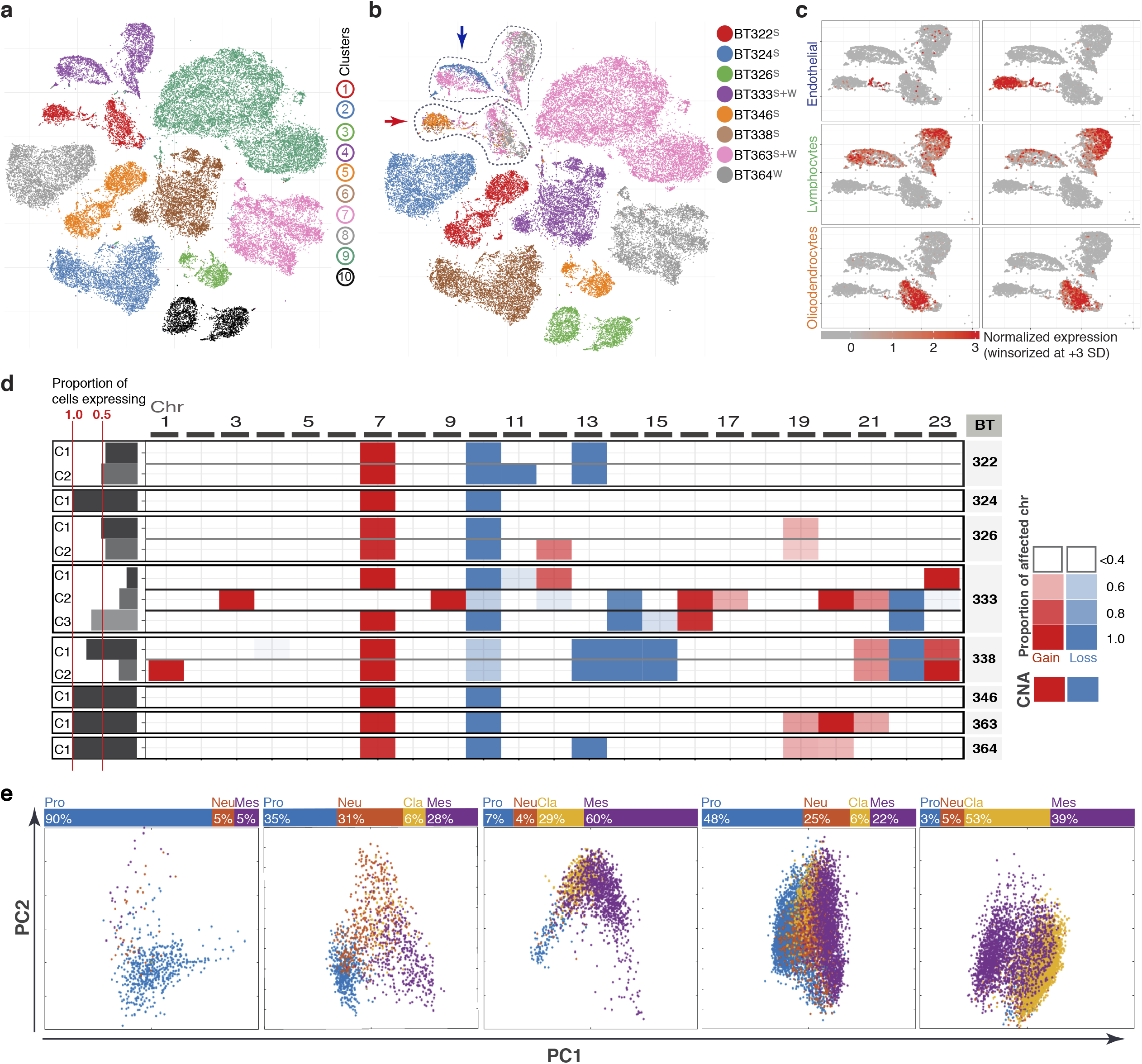
Single-cell RNA sequencing highlights genomic and transcriptomic heterogeneity in glioblastoma. **(a)** t-distributed stochastic neighbour embedding (tSNE) of location-averaged transcriptome for all tumour cells (whole tumour and glioma stem cells) from 8 patients. Cells are colored by cluster. S corresponds to glioma stem cell samples, W corresponds to whole samples **(b)** tSNE of location-averaged transcriptome for all tumour cells colored by patient. Cancer cells cluster by patient, whereas normal cells from all patients cluster together (encircled clusters indicated by arrows). **(c)** Expression of endothelial (CD34 and ESAM), lymphocyte (CD53 and CD74) and oligodendrocyte (MOG and MBP) genes in clusters devoid of CNAs. **(d)** Copy number aberrations heatmap for all main clones of each patient. The transparency shows how much of the chromosome is affected, starting at 50%. All findings shown were significant (p<0.001) using the Wilcoxon test. **(e)** Cell-cycle corrected transcriptome principal component analysis of whole tumour cells for each patient. Cells are colored according to their TCGA subtype. Most TCGA subtypes were detected in each patient.

Occasionally, cells from a given patient generated two or three cancer groupings by t-distributed stochastic neighbour embedding (tSNE), indicating different clones within a tumour (Fig. 1a). To better characterize these clones, we pooled cells from the cancer clusters of each tumour and reclustered them with our location-averaged data. We determined the correct number of clusters by finding the most stable solution (Supplementary Fig. 1c). We detected one to three clones for each tumour. These clones differed by a limited number of CNAs (Fig. 1d). Together, these findings demonstrate intertumoural and intratumoural genomic heterogeneity.

We then assessed intratumoural heterogeneity in the whole tumours based on single-cell transcriptomic data. To reduce noise, we selected genes with high variance in each sample. We performed principal components analysis (PCA) one sample at a time to avoid interpatient heterogeneity. PCA analysis finds the combination of genes best able to explain the transcriptomic variation between cells in a sample. At this point, we observed variations in gene expression due to the cell cycle (Supplementary Fig. 1e). To reduce cell cycle effect and reveal other sources of heterogeneity, we scored each cell for the various phases of the cell cycle ^31, 32^ and chose cells scoring less than or equal to 0 in both the G2/M and G1/S scores (defined as the non-cycling cells) as input to calculate the PCA eigenvectors. The full dataset (cycling and non-cycling cells) was then projected onto these eigenvectors for visualization.

Once the cell cycle effect was removed, variability in gene expression profiles remained apparent within tumours and between tumours. We identified three to four TCGA subtypes in each tumour, as was previously shown ^12^ (Fig. 1e). Importantly, cells with different TCGA subtypes were often separated by the first of second principal components (PCs), indicating that these subtypes accurately describe a portion of the intrinsic heterogeneity of each tumour. Also, in each tumour, cells with different TCGA subtypes did not necessarily belong to different CNA clones (Supplementary Fig. 2a); however, different proportions of TCGA subtypes were observed between some clones within individual tumours. This is consistent with results from the TCGA indicating that genomic aberrations do not perfectly predict a subtype.

Some trends were observed from this whole tumour analysis. Cells expressing neuronal genes such as STMN4 and SOX11 were not found in cells expressing astrocytic genes such as AQP4 and APOE, and these cells were separated by the first or second PC in each tumour (Supplementary Fig. 2b). Also, stemness genes such as ASCL1 and OLIG2 were often highly expressed in cells with PC values intermediate to cells expressing neuronal and astrocytic genes (Supplementary Fig. 2b). These findings suggest organization within heterogeneity.

### Glioblastoma stem cells develop following a conserved neurodevelopmental hierarchy

We then applied the same PCA analysis strategy to assess transcriptomic heterogeneity in enriched GSCs from each tumour. The cycling-free PCA strategy described above was used since not all cells were cycling (Supplementary Fig. 2c).

For each GSC enriched tumour sample, we found that the first PC separates cells into neural developmental lineages. GSCs expressing neuronal genes such as CD24, SOX11 and DCX were mutually exclusive from cells expressing astrocytic genes such as GFAP, APOE, AQP4, CD44, CD9, and VIM (Fig. 2a). To assess the conservation of these gene programs across patients, we ranked genes by strength of influence on PC1 and found a strong correlation of these ranks between samples (R-squared = 0.77, Fig. 2b). GSCs with intermediate PC1 values express progenitor cell genes such as SOX4, OLIG2, and ASCL1 (Fig. 2a). In some samples, these cells had high PC2 values; however, this was not apparent in all samples and the rank correlation was lower (R-squared = 0.34, Supplementary Fig. 2d). We validated the differential gene expression profiles of enriched GSCs and whole tumour cell populations using flow cytometry. In general, cells do not coexpress neuronal (e.g., CD24) and astrocytic (e.g., CD44) markers (Fig. 2c). Together, these data suggest that GSCs are organized into progenitor, neuronal, and astrocytic gene expression programs, resembling a developing brain.

**Figure 2.**
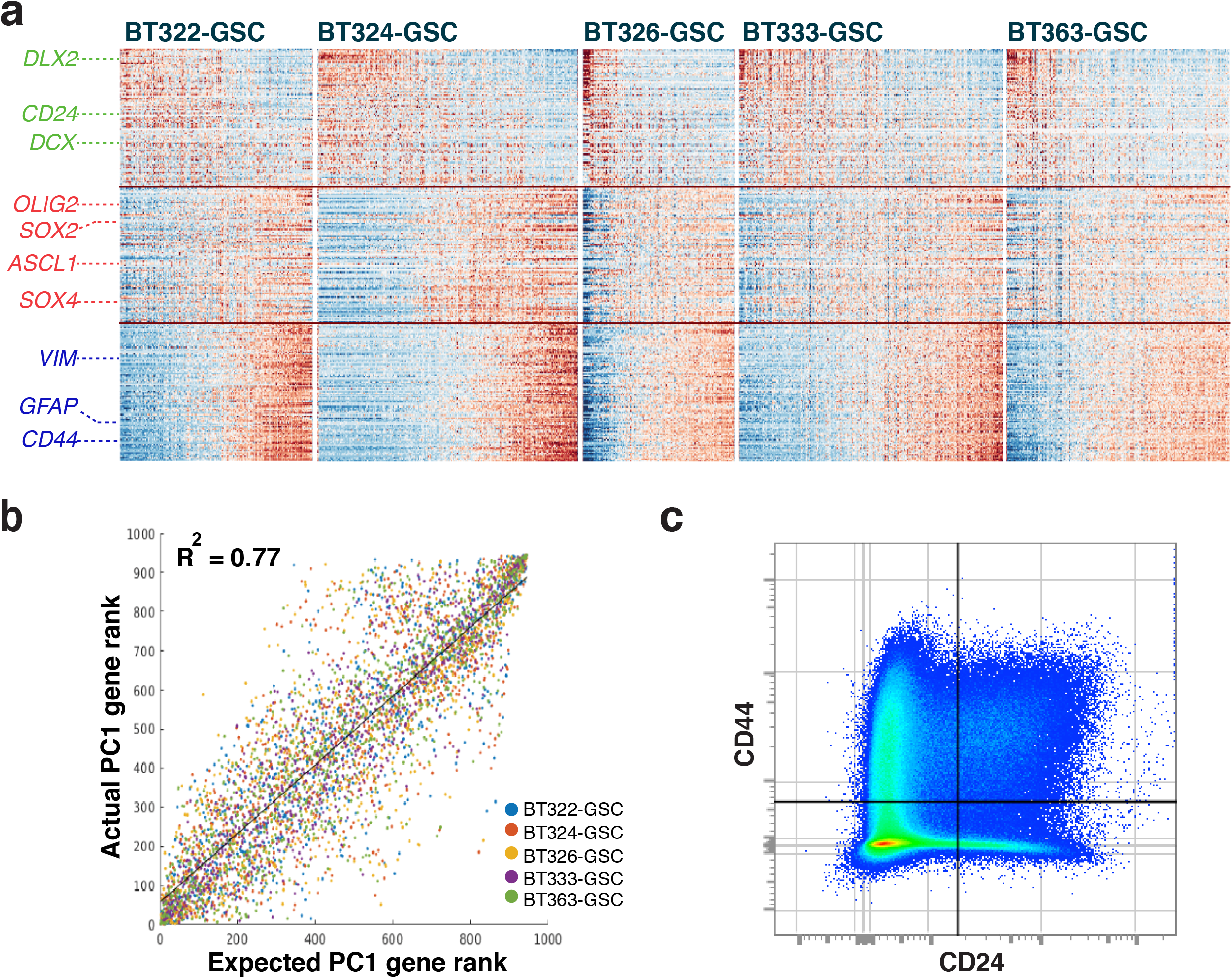
Glioblastoma stem cells develop following a conserved neurodevelopmental hierarchy. **(a)** Enriched-glioblastoma stem cell (GSC) gene expression heatmaps showing relative gene expression (raw data) correlated with PC1 per patient. These maps are separated into 3 rows: top row - 100 genes with the lowest value for PC1 loading; bottom row - 100 genes with the highest value for PC1 loading; middle row - 100 genes with the highest value for PC2 loading. These gene signatures correspond to neuronal, astrocytic and progenitor signatures, respectively. **(b)** Expected and actual rank of genes by PC1 correlation. The actual gene rank (y-axis, one point per sample) correlates strongly with the expected gene rank (x-axis) in all patients. **(c)** Flow cytometry analysis of GSCs and whole tumour demonstrating mutually exclusive expression of CD24 and CD44.

### Single-cell RNA sequencing of the developing brain and the identification of glial progenitor cells

If glioblastoma is organized into programs reflecting normal brain development, then a direct comparison to the developing brain at a single cell level should provide additional insight. We performed scRNAseq on freshly isolated cells from the telencephalon of four human fetuses ranging from 13 to 21 weeks of gestation. Fluorescence-assisted cell sorting (FACS) was used to remove most microglia (CD45-positive) and endothelial cells (CD31-positive) from the samples, and to select CD133-positive cells in order to improve the resolution of progenitor and neural stem cell populations ^33^. By sequencing both the total and the CD133-positive cell populations, we aimed to maintain cellular representation of development. We sequenced 12 544 cells from the total unsorted population, and 10 093 cells from the CD133-positive population.

Total and CD133-positive datasets from all fetal brains were combined in silico (Supplementary Fig. 3a) after excluding ependymal cells (see methods), and the Louvain community detection algorithm was used to group cells into cell types (Fig. 3a, b). By varying the resolution parameter of the algorithm, we chose the most stable clustering solution (Fig. 3b and Supplementary 3b). Two modifications were made to this solution. The first was to consolidate excitatory neurons-four clusters coincided on the tSNE plot and strongly expressed neuronal genes such as NEUROD6, SYT1, and STMN2. Second, a smaller cluster spanned multiple apparent groups on the tSNE plot. Differing expression of OLIG2, PDGFRA, GFAP, APOE, ASCL1, and AQP4, amongst other genes, were apparent within this cluster (Fig. 3c and Table 1). We thus opted to use the algorithm described above on this group of cells, which further separated it into three clusters consistent with: oligodendrocyte lineage cells (OLCs); astrocytes; and a previously unidentified glial cell type. This generated a total of ten cell clusters (Fig. 3a). Differential gene expression analysis of these clusters (Table 1) identified cell types spanning all of the main cell lineages previously identified in similar scRNAseq studies ^34, 35^. CD133-positive cells were found in all clusters/cell types, but were enriched in the radial glia, neuronal progenitors, and committed glial cell clusters (Fig. 3d and Supplementary Fig. 3c).

**Figure 3.**
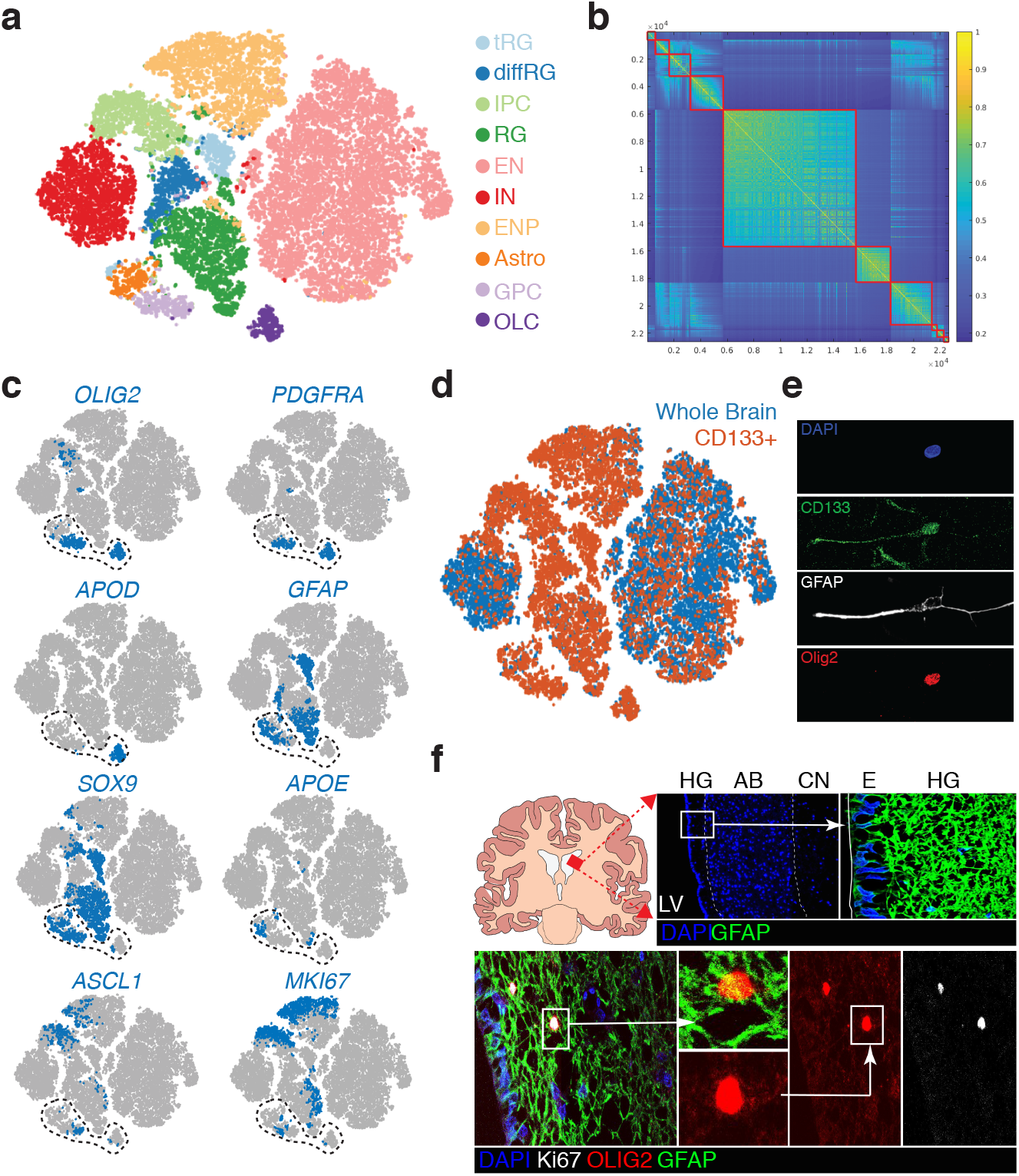
Single-cell RNA sequencing of the developing brain and the identification of glial progenitor cells. **(a)** T-distributed stochastic neighbour embedding (tSNE) map of human fetal brain cells by cluster or cell type. Datasets from total cells and CD133-positive cells were combined. Cells are colored by cell type. tRG – truncated radial glia, diffRG – differentiating radial glia, IPC – inhibitory neuronal progenitor, RG – radial glia, EN – excitatory neuron, IN – inhibitory neuron, ENP – excitatory neuronal progenitor, Astro – astrocyte, GPC – glial progenitor cell, OLC – oligo-lineage cells. **(b)** Similarity matrix of fetal cells ordered by cluster. **(c)** tSNE maps of human fetal brain cells showing cell type expression of OLIG2, PDGFRA, GFAP, ASCL1, SOX9, MKI67. Expression is averaged to the twenty closest neighbours in principal component (PC) space. **(d)** tSNE map of total human fetal brain cells and CD133-positive fetal brain cells. **(e)** Representative example of a freshly cultured fetal neural stem cell coexpressing CD133, OLIG2, and GFAP. **(f)** Immunofluorescence analysis of the adult human subventricular zone (SVZ). Top row, schematic and anatomic structure of the SVZ. HG – hypocellular gap, AB – astrocytic band, E – ependymal cells, LV – lateral ventricle, CN – caudate nucleus. Bottom row, identification of dividing cells with marker expression corresponding to glial progenitor cells

Two CD133-positive cell types did not fit with previously identified gene signatures ^34–36^. The first was detected mainly in the 21-week brain and highly expressed genes such as VIM, GFAP, OLIG1, GLI3, and EOMES. We have tentatively termed them differentiating radial glia cells due to their mixed signature (Fig. 3a). The second cell type, the unidentified glial cell cluster mentioned above, was detected at all gestational ages and strongly expressed oligodendrocyte lineage genes (e.g., OLIG1, OLIG2, and PDGFRA), glial/astrocytic lineage genes (e.g., GFAP, SOX9, HOPX, HEPACAM, and VIM), and progenitor genes (e.g., ASCL1, MKI67, and HES6) (Fig. 3c and Table 1). However, it did not express differentation markers found in astrocytes or OLCs such as APOE and APOD, respectively (Fig. 3c). It also lacked the high gene complexity and UMI counts seen in doublets (Supplementary Fig. 3d). This mixed gene signature is compatible with that of a bipotential glial progenitor cell (GPC). Notably, this GPC signature was almost exclusively identified in CD133-sorted cells (Fig. 3a, d and Supplementary Fig. 3c), which likely explains why it was not previously detected ^34, 35^. The existence of cells expressing these GPC markers was confirmed in first passage culture of fetal brain cells derived from one of the fetal brains sequenced (Fig. 3e), and in the subventricular zone of the adult human brain (Fig. 3f).

### Creation of a fetal brain roadmap to uncover the organization of glioblastoma

Upon developing a fully indexed dataset of the developing human brain, we aimed to parallel each cancer cell to a fetal brain cell type. To do so, we developed a roadmap technique that enables the projection of every cancer cell onto the fetal dataset, similar to the shadow of an object cast onto a nearby surface.

To build the final roadmap, we first found the fetal cell types which best represent the cancer. This was accomplished by determining which fetal brain cell type was nearest to, or captured, each cancer cell. Ninety-four percent of whole tumour cells were captured by five fetal brain cell types: neurons; astrocytes; OLCs; truncated radial glia (tRG); and GPCs. Sixty-seven percent of whole tumour cells were captured by either astrocytes, OLCs, or GPCs (Supplementary Fig. 4a). The proportion of enriched GSCs captured by GPCs was substantially greater than that of whole tumour cells (Supplementary Fig. 4a, b). Some neuronal cell types also captured cancer cells, as predicted by the GSC data (Fig. 2a). Surprisingly, interneurons captured more cells than excitatory neurons.

A limitation of this fetal dataset is the absence of a proper correlate to mesenchymal cancer cells. As such, they were inappropriately captured by tRGs. We believe this mapping is incorrect because the tRG signature does not match that of mesenchymal cancer cells. These cancer cells lack expression of common tRGs genes such as AQP4, FAM107A, SOX9, and GLI3. Also, only 1.1% of enriched GSCs are captured by tRGs (Supplementary Fig. 4b), suggesting these mesenchymal cancer cells are more differentiated than tRGs. Consequently, four cell types were used to construct the roadmap: astrocytes; GPCs; OLCs; and interneurons.

The final roadmap needs to not only capture heterogeneity between cancer cell types, but also heterogeneity within a given cancer cell type to accurately represent the continuous process of differentiation. Therefore, we used PCA on an equal number of fetal astrocytes, GPCs, OLCs, and interneurons. This fetal PC space acts as the roadmap. We then used diffusion embedding ^37, 38^ to better represent the differentiation process in 3D. In this diffusion roadmap, GPCs are found at the junction of the oligodendrocytic, astrocytic, and neuronal lineages (Fig. 4a).

**Figure 4.**
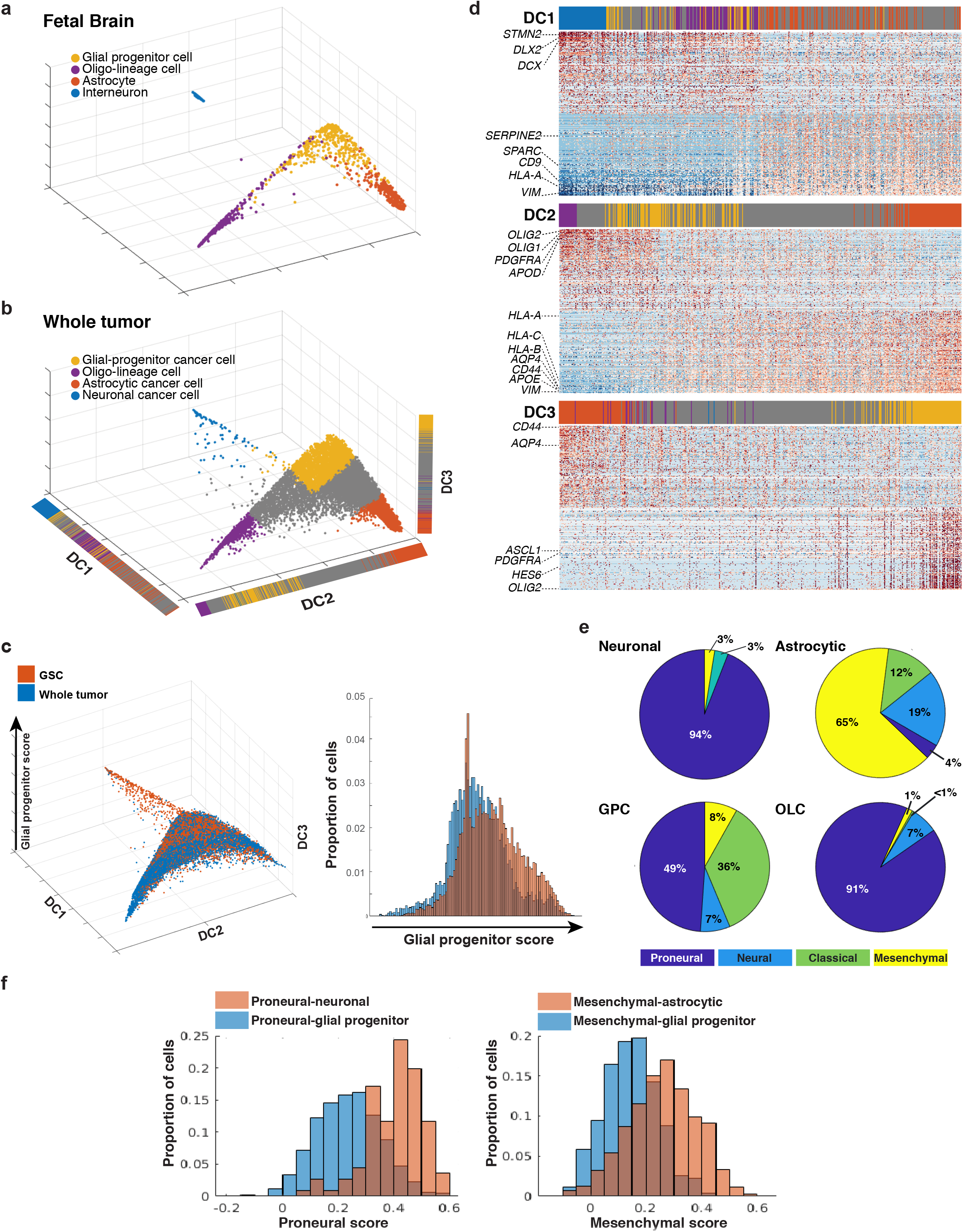
Fetal brain roadmap reveals a glioblastoma trilineage hierarchy centered on progenitor cancer cells. **(a)** Diffusion plot of the projection of select fetal cell types onto the roadmap. Cells are colored by the cell type they were attributed in Fig. 3a. **(b)** Diffusion plot of the projection of all whole tumour cells onto the roadmap. Cells are colored based on their classification by linear discriminant analysis (LDA). Unclassified cells were coloured grey. **(c)** Diffusion plot showing the location of glioma stem cells (GSCs) relative to whole tumour cells (left) and histogram of glial progenitor score for GSCs and whole tumour cells (right). An increase in proportion of cells with higher glial progenitor scores is seen in GSCs (p< 1e-21). **(d)** Heatmaps showing relative gene expression (raw data) for cells ordered by each of the diffusion components of the roadmap. Genes are ordered from most correlated to least correlated with the diffusion component. Only the 150 most and 150 least correlated genes are shown. Top color bar indicates cell type classification from the LDA. Each color corresponds to the same classification as in b. **(e)** Pie chart for TCGA subtype by cell type. Cell types are based on the LDA classification for all whole tumour cells. **(f)** Histograms comparing the TCGA subtype score between whole tumour cells of a given cell type and glial progenitor cancer cells of the same TCGA subtype. For cells of a given TCGA subtype, glial progenitor cancer cells consistently show lower TCGA score in that subtype than the more differentiated cancer cells of the cell type, which makes up the majority of cells with this TCGA subtype. GPC – glial progenitor cancer cell, Mes – mesenchymal, OLC – oligo-lin-eage cell.

### Fetal brain roadmap reveals a glioblastoma trilineage hierarchy centered on progenitor cancer cells

We projected all cancer cells onto this roadmap and used the first three components of diffusion embedding as each cell’s coordinate in the hierarchy (Fig. 4b). GSCs and whole tumour cells from all patients overlapped (Fig. 4c) despite significant variations in lineage proportions between patients in the whole tumour samples (Supplementary Fig. 4c). The GPC signature was the only one robustly expressed in all patients. To visualize gene signatures, we ordered cancer cells according to all three diffusion components (DCs) individually and found genes that correlated most with this order (Fig. 4d and Table 2). Cancer cells expressing an OLC signature (e.g., OLIG1, APOD; Supplementary Fig. 4d) or an astrocytic signature (e.g., GFAP, AQP4) were found at either end of DC2; cancer cells expressing a neuronal signature (e.g., STMN2, DLX2) were found at the end of DC1; and cancer cells expressing a GPC signature (e.g., OLIG2, ASCL1, HES6) were found at the end of DC3. We therefore defined DC3 as the glial progenitor score. This organization reveals a glial progenitor-centered trilineage hierarchical organization of whole tumour and enriched GSCs.

When comparing enriched GSCs and whole tumour cells, we found a significant shift of GSCs towards higher values on the glial progenitor score (p<1E-21), and a shift towards intermediate values of DC2 (Fig. 4c and Supplementary Fig. 4e). The proportion of OLCs in enriched GSCs and whole tumour was the same (Supplementary Fig. 4e). These data show that glial progenitor cancer cells are enriched in GSC culture conditions, and the number of more differentiated astrocytic cancer cells is decreased in stem cell culture conditions.

Lastly, cancer cells from whole tumour were classified into cell types using a linear discriminant analysis (LDA) with the fetal cells as a training set (Fig. 4a). Cells that could not be classified with a probability of error less than 0.01% were left unclassified (Fig. 4b); these correspond to cells with intermediate signatures. We compared the TCGA subtypes according to the classified cell types (Fig. 4e). As predicted by previous work ^19^, neuronal and oligo-lineage cancer cells were almost exclusively proneural. Astrocytic cancer cells were mesenchymal, classical, or neural. Finally, glial progenitor cancer cells were mostly proneural (Fig 4e), but some classical and mesenchymal type cells were also found. Interestingly, we found that the mesenchymal type glial progenitor cancer cells had lower mesenchymal scores than the mesenchymal astrocytic cancer cells (Fig. 4f, p<1E-22). Similarly, proneural glial progenitor cancer cells had lower proneural scores than the proneural neuronal cancer cells (p< 1E-22). This is in keeping with a hierarchical model of cancer differentiation in which progenitors can lean in one direction for differentiation, but do not express the differentiated gene program as strongly as the more differentiated cancer cells.

### Progenitor cancer cells are the most proliferative cancer cells

We next set out to determine which population of cancer cells is the most proliferative, with the goal of developing therapeutics to target this cell population. Based on the expression of cell cycle genes, we defined a cycling cell as one with a G1/S or G2/M score greater than 1.5, as was done previously ^32^. We then calculated the proportion of cycling cells as a function of their glial progenitor score. We found almost all cycling cancer cells had high glial progenitor scores (Fig. 5a, b).

**Figure 5.**
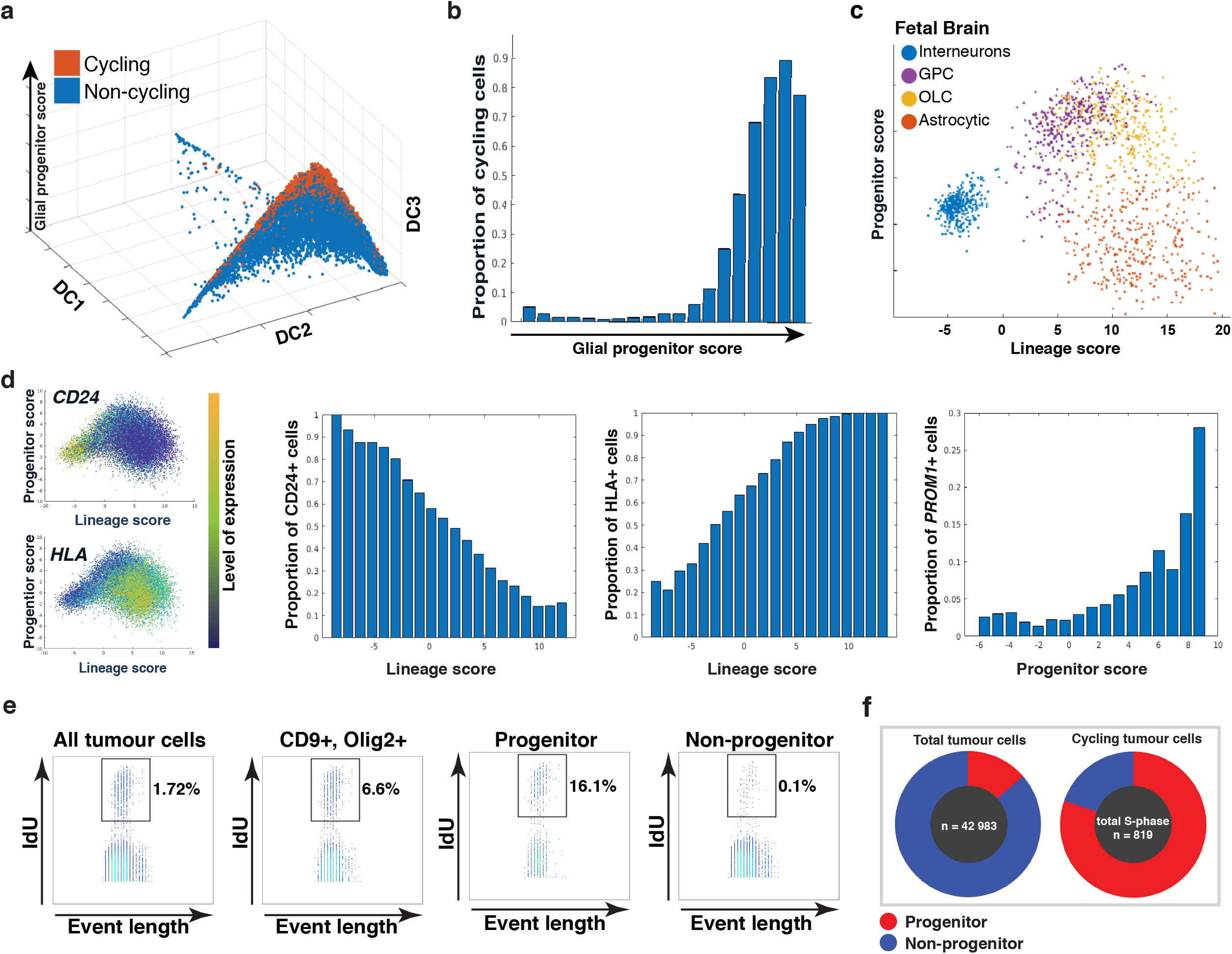
Progenitor cancer cells are the most proliferative cancer cells. **(a)** Diffusion plot of the roadmap of whole tumour cancer cells showing that cycling cells are predominantly glial progenitor cancer cells. Cycling cells are defined by >1.5 in either the G1/S or G2/M scores. **(b)** Bar chart showing that the proportion of cycling cells increases with increasing glial progenitor score. **(c)** Simplified roadmap in principal component space with select fetal cell types. Projected OLCs and GPCs overlap and are high for progenitor score, while interneurons and astrocytes are lower in progenitor score, but occupy opposite ends of the lineage score. **(d)** Projection of glioma stem cells (GSCs) on the simplified roadmap highlights the location of CD24 and HLA within the hierarchy. For each gene, the simplified roadmap projection shows the expression of this gene in GSCs, and the histograms show the proportion of cells where CD24, HLA and PROM1(CD133) were detected at differing positions in hierarchy. **(e)** Mass cytometry pseudo-color dot plots showing the proportion of whole tumour cells, progenitor cancer cells, and non-progenitor cells that are in S-phase. The progenitor cancer cell population has the highest proportion of cells in S-phase. **(f)** Mass cytometry showing that progenitor cancer cells are the main cycling cell population in the tumour. Pie charts showing the proportion of progenitor cells (CD133+, OLIG2+, PDGFRA+) in the tumour (left) and the cycling population (right).

We then aimed to validate this result using single-cell proteomic analysis. To do so, we generated protein marker panels representative of each cancer cell type. Since the OLCs and glial progenitor cancer cell types are both progenitor in nature (OLCs contain oligo-progenitor cells) and a reliable discriminant surface marker could not be found, we considered these two cancer cell types together as progenitors. This was achieved by adding the two roadmap PCs specific to each of these two fetal cell types – we called this the progenitor score (Fig. 5c). The remaining PC mainly differentiated fetal astrocytes from fetal interneurons – we called this the lineage score. This new simplified roadmap therefore had a progenitor cancer cell type at the apex of two lineages (Fig. 5c), similar to the GSC PCA. We projected GSCs onto this modified roadmap and selected genes encoding cell surface protein markers which most strongly correlated with the lineage scores (Fig. 5d and Table 3). For the purposes of cytometry assays and sorting, we defined CD24^+^/CD133^−^/ CD9^−^ as neuronal cancer cells, CD9^+^/CD44^+^/CD133^−^ as astrocytic cancer cells, and PDGFRA^+^/CD133^+^ as progenitor cancer cells. For the mass cytometry assay, OLIG2 was included as an intracellular marker for progenitor cancer cells.

Using the progenitor cancer cell marker panel, and a validated cell cycle marker panel ^39, 40^, we analyzed 37 185 cancer cells, and found 640 cells in S-phase (1.7%) (Fig. 5e). We found that 16.1% of the progenitor cancer cell population was in the S-phase. These progenitor cancer cells make up only 14% of the total tumour population yet account for 80% of all cycling cells (Fig. 5f). Interestingly, most of the remaining cycling cells express a subset of the progenitor signature, highlighting the continuous nature of differentiation. In contrast, 0.1% of cells without progenitor markers were found to be cycling (Fig. 5e), while accounting for 77.3% of the total tumour population. Similarly, tumour immunolabeling, using Ki67 as a marker of cell proliferation, showed that the percentage of cycling cells in the CD133-positive population is significantly higher than that of CD133-negative population in two patients (Supplementary Fig. 5a).

### Progenitor cancer cells are drivers of chemoresistance and tumour growth

Resistance to conventional chemotherapies and tumourigenicity are hallmarks of CSCs ^6, 17, 18^. These properties, however, are derived from studies that have considered the CSC compartment to be uniform, not one displaying heterogeneity driven by a hierarchical developmental organization. To evaluate GSC chemoresistance and tumourigenicity considering hierarchy and lineage, we sorted them into three types – progenitor, neuronal, and astrocytic, based on gene expression signatures described above.

Three patient-derived GSC lines were separated into these types and treated with temozolomide (TMZ), the standard of care glioblastoma chemotherapy. Variable doses were required to achieve responses in different cell lines, correlating with the methylguanine methyltransferase status of the tumour. We found that progenitor GSCs either did not respond or responded less to TMZ than the more differentiated GSCs. (Fig. 6a and Supplementary Fig. 5b).

**Figure 6.**
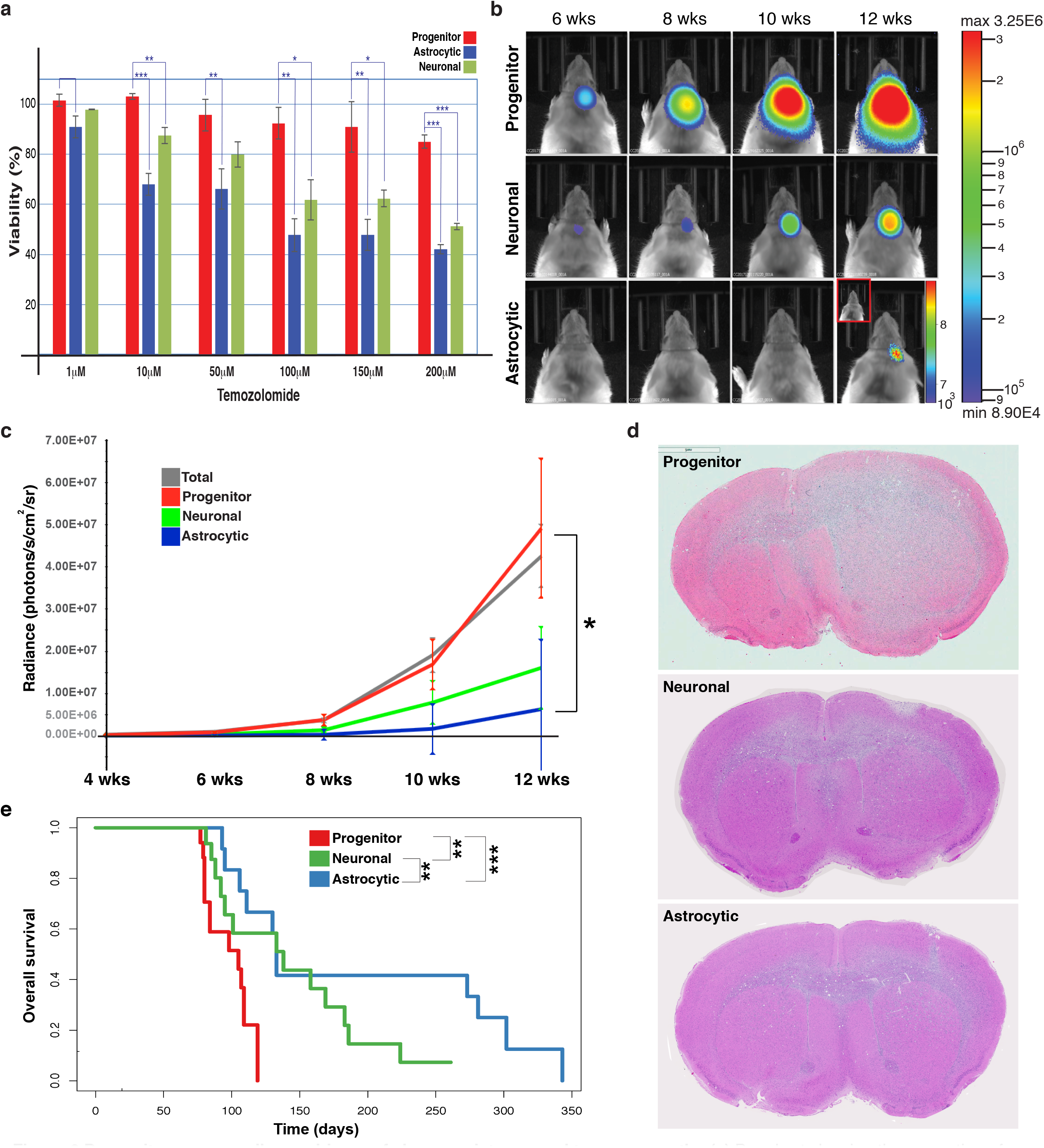
Progenitor cancer cells are drivers of chemoresistance and tumour growth. **(a)** Bar chart showing the proportion of viable glioma stem cells (GSCs, patient line BT390) sorted by type following 5 days of temozolomide (TMZ) treatment, normalized to corresponding vehicle control. See Supplementary Fig. 5b for additional patients. **(b)** Select bioluminescence images from mice implanted with GSCs sorted by type. Mice implanted with progenitor GSCs exhibit a more rapid tumour growth compared to those implanted with neuronal or astrocytic GSCs. **(c)** Average bioluminescence intensity over time for mice xenografts injected with different GSC types for BT333 (24 mice). Data are represented as mean +/- SE. * indicates p-value less than 0.05. **(d)** A representative mouse from each GSC group was sacrificed at 12 weeks and the corresponding H&E stained images are shown. **(e)** Kaplan-Meier survival curves for mice implanted with different GSC types (n=47). Univariate Cox proportional Hazard Model shows a significant difference in survival between progenitor GSC and neuronal or astrocytic GSC xenografts. *** indicates p-values less than 0.001, ** indicates p-values less than 0.01, * indicates p-value less than 0.05.

We then assessed the influence of hierarchy and lineage on tumour forming capacity. Forty-seven mice were orthotopically xenografted with progenitor, neuronal, astrocytic, or total GSCs from 3 different patients in near-limiting dilution. Bioluminescence was assessed over 12 weeks, and survival was determined. We observed earlier tumour formation, and a more rapid increase in tumour signal, for all mice implanted with progenitor or total GSCs. In mice implanted with astrocytic or neuronal GSCs the tumour formation and signal increase was either absent or significantly delayed by up to 3 months (Fig. 6b-d). Consistently, mice implanted with progenitor GSCs had a significantly lower survival time than those implanted with neuronal (OR 0.26, p<0.01) or astrocytic (OR 0.05, p<0.001) GSCs (Fig. 6e).

Together, these results identify a hierarchy of tumourigenicity and chemoresistance in GSCs, with progenitor cancer cells being the most chemoresistant and tumourigenic.

### Pathways enriched in progenitor cancer cells expose therapeutic opportunities

Since progenitor GSCs are the most chemoresistant and tumourigenic cancer cell population, we aimed to leverage our hierarchy and transcriptomic data to find targets relevant to this cancer cell population.

We used the LDA classification of whole tumour cells described above to separate cells into cell types. We selected the GPC and astrocytic groups for the analysis to specifically compare the progenitor population to the most abundant cell type in the cancer. We performed gene set enrichment analysis (GSEA) in a manner similar to previously described methodologies ^41^. Briefly, each gene in the dataset was ranked for its correlation to progenitor cancer cells versus astrocytic cancer cells. For each gene set ^41–43^, the Mann-Whitney U statistic was used to determine if genes within the set had higher correlation with the progenitor cancer cells than genes not present in the set. After correcting for multiple testing, we identified pathways with a significant enrichment in progenitor cancer cells (Table 4). Hits with significant and strong correlations were found in pathways such as EZH2, FOXM1, and Wnt, previously established pathways relevant to cancer stem cell self-renewal and tumourigenicity ^44–48^.

Pathways of previously unknown significance in GSCs were also detected. Of these, the E2F pathway was the most significant, and it was thus selected to test our target identification method. The E2F gene family regulates cell cycle and is important for progenitor cell survival ^49^. The E2F gene set involves many of the regulating targets of the transcription factor E2F; therefore, E2F inhibition was selected to target this pathway. HLM006474 is a small molecule inhibitor that prevents E2F4 binding to DNA. It has been shown to cause senescence of gastric cancer cells ^50^, and to reduce proliferation and survival of melanocytic cells and lung cancer cells *in vitro* ^51, 52^. E2F4 expression in glioblastoma tissue has been shown ^53^. To our knowledge, our work provides the first description of its importance in GSCs.

We tested the effect of E2F inhibition in progenitor, neuronal, and astrocytic GSCs following HLM006474 treatment. Proliferation and survival of progenitor GSCs was significantly reduced compared to neuronal and astrocytic GSCs (Fig. 7a). This differential sensitivity was also observed in a sphere forming capacity assay (Fig. 7b, c) and serum-free vs serum-differentiated GSCs (Supplementary Fig. 6a). On target E2F inhibition was confirmed^51^ (Supplementary Fig. 6b, c). Together, these data show that targeting E2F preferentially affects progenitor GSC proliferation.

**Figure 7.**
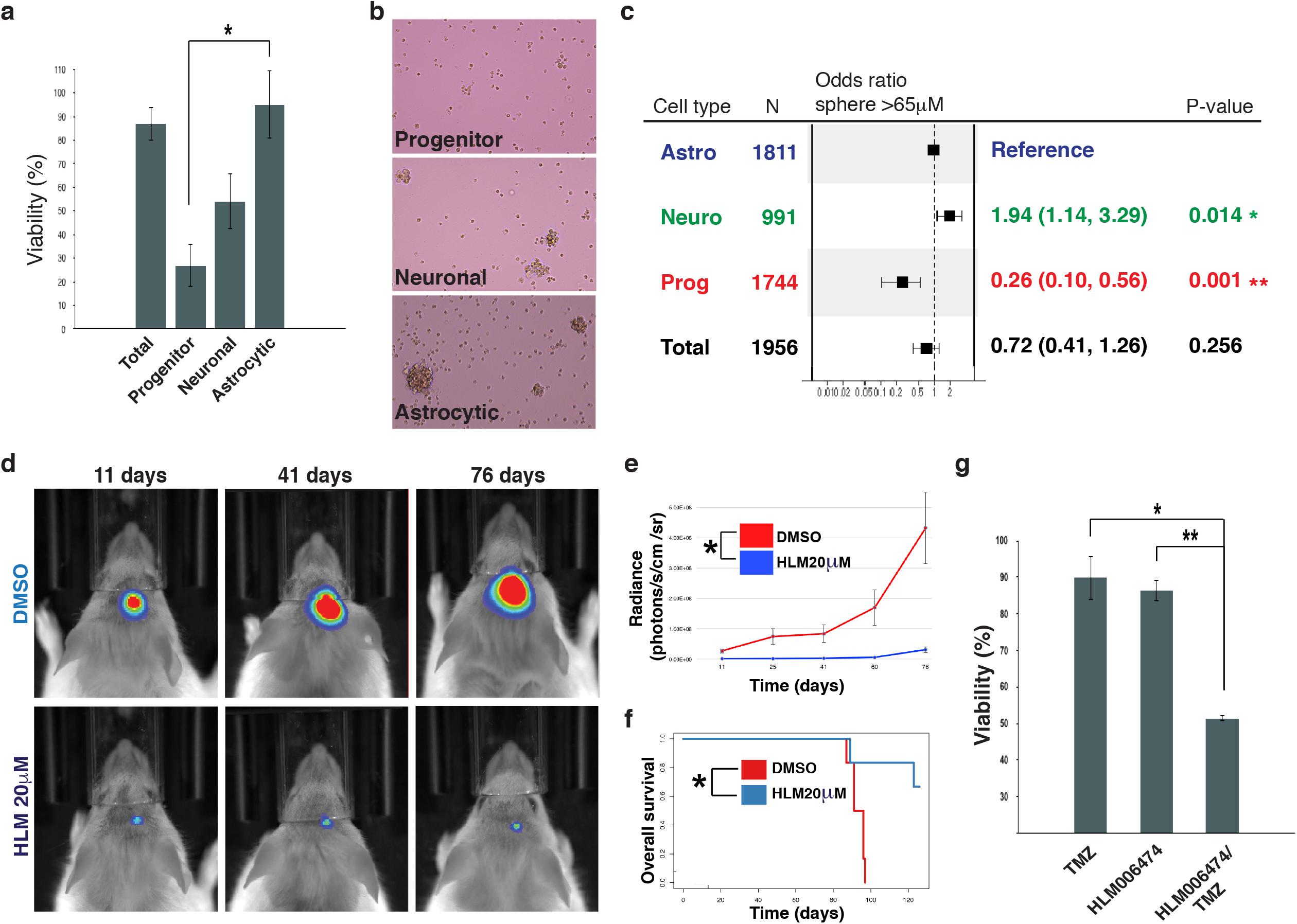
Pathways enriched in progenitor cancer cells expose therapeutic opportunities. **(a)** Bar chart showing the proportion of viable glioma stem cells (GSCs) sorted by type followed by 7 days of HLM006474 treatment, normalized to corresponding vehicle control. Data are represented as mean +/- SE. * indicates p<0.05. **(b)** Representative images of GSCs sorted by type and treated in HLM006474 for 7 days (images correspond to 7a). **(c)** Forest plot showing the odds ratio of forming a tumour sphere greater than 65μm following 7 days of HLM006474 treatment, calculated using a multivariate logistic regression with the astrocytic GSC type as a reference, controlled for patient cell line. There was no significant difference between the 2 GSC lines (p>0.2), 95% confidence intervals are shown. * indicates p-value less than 0.05, ** p<0.01. **(d)** Bioluminescence images and **(e)** signals from representative mice treated with 20μM HLM006474 vs DMSO with corresponding **(f)** Kaplan-Meier survival plot (n=16, 8 per group). Data are represented as mean +/- SE. * indicates p-value less than 0.05, ** p<0.01. **(g)** Bar chart showing the proportion of viable GSCs sorted by type followed by 6 days of TMZ treatment, 6 days of HLM006474 treatment, and 3 days of TMZ treatment followed by 3 days of HLM006474 treatment, normalized to corresponding vehicle control.

We tested the effects of E2F4 inhibition *in vivo*. Pooled GSCs, treated with HLM006474 or vehicle for 3 days, were orthotopically xenografted. A significant reduction in tumour growth (Fig. 7d, e), and improved survival (Fig. 7f, p-value = 0.03) in the HLM006474 treated mice was observed.

Since E2F inhibition is effective in progenitor GSCs, and TMZ chemotherapy is more effective in differentiated cells ^18^, we reasoned that HLM006474 combined with TMZ would be a more effective treatment for the total GSC compartment than each individually. We sequentially treated GSCs with HLM006474 followed by TMZ chemotherapy at TMZ doses that are ineffective as monotherapy. We observed a significant decrease in proliferation and cell survival using this combination therapy compared to monotherapy (Fig. 7g).

## Discussion

Intratumoural and interpatient heterogeneity are hallmarks of many cancers ^1, 2, 4^. Heterogeneity complicates efforts to understand cancer development, progression, and the development of effective therapeutics. Here, we show that the normal developing human brain can be used as a roadmap to elucidate brain cancer development, and, in so doing, reveal that glioblastoma develops along conserved neurodevelopmental gene programs shedding new light on the cancer stem cell hierarchy and the origins of heterogeneity.

Recently, scRNAseq characterization of human fetal brain cells described the transcriptomic signature of many cell types within the developing brain ^34, 35^. By increasing the number of cells sequenced, and enriching for neural stem cells, we uncovered a previously unidentified cell type with a transcriptomic signature suggestive of a GPC. Additional work such as fate mapping will be necessary to uncover the exact position of these cells within the developmental hierarchy of the brain. This finding sheds new light on gliogenesis, and highlights the role of cell sorting to detect rare cell populations.

Using this developmental data as a roadmap, we show that isocitrate dehydrogenase (IDH) wild-type glioblastoma is organized into three lineages that correspond to all three normal neural lineages: astrocytic; neuronal; and oligodendrocytic. We also found a cluster of cells that exists at the intersection of these lineages and corresponds transcriptomically to progenitors and functionally to cancer stem cells. The closest transcriptomic parallel of this cell cluster in the normal developing human brain are GPCs. We propose that the cell of origin of glioblastoma, whether a GPC/OPC or another cell type nearby in the brain stem cell hierarchy, also possesses such pluripotency. A genetic mouse model studying glioma origin suggested that OPCs are candidate cells of origin ^54^. These cells expressed PDGFRα, Olig2, and occasionally nestin. In our dataset, both GPCs and OPCs express PDGFRA and OLIG2, but NESTIN expression is restricted to GPCs. In contrast, scRNAseq studies of IDH mutant gliomas and H3K27M pediatric gliomas identified only two lineages, astrocytic and oligodendrocytic ^30–32^. This suggests a different cell of origin in these pathologies than in glioblastoma and may underlie the disparate natural histories and treatment responses between these cancer types.

Our data also show that progenitor cancer cells are the cancer cell type with the highest rates of proliferation, more so than differentiated cancer cells. As genetic anomalies are most often acquired during the cell cycle, it is likely that new clones arise within this progenitor population and propagate down the lineages as their progeny differentiate. Consistently, we found that heterogeneity develops very early in cancer development, within progenitor cancer cells. In fact, the heterogeneity that develops within glial progenitor cancer cells is sufficient to generate all four TCGA subtypes within this cell cluster. These findings are supported by Bhat et al. ^55^ and Mao et al.^56^ who showed that GSCs can express a proneural or mesenchymal profile. We suspect that specific genomic anomalies skew differentiation towards one lineage or another, giving rise to the observed TCGA subtypes.

Our findings show that progenitor cancer cells give rise to the transcriptomic heterogeneity within the tumour, are the most rapidly cycling cancer cell type, and are the most tumourigenic in xenograft models, more so than distal cancer stem cells or differentiated cancer cells. Together, these findings are relevant to cancer biology and therapeutics development. These rapidly cycling progenitor cancer cells are the earliest detectable cancer cells in the hierarchy and thus serve as a prime cell population to target. Identification of mutations or driver events within this cell population, or the identification of signaling pathway alterations between progenitor cancer cells and more differentiated cancer cells will likely yield meaningful new therapeutic targets.

To that end, we leveraged our transcriptomic data and conserved hierarchical neurodevelopmental classification to identify therapeutic targets relevant to progenitor cancer cells in all patients. HLM006474, an E2F blocker, shows pronounced activity towards progenitor GSCs versus GSCs that have differentiated towards the neuronal or astrocytic lineages. We showed that E2F4 inhibition significantly hampered tumour growth *in vivo*. Pre-clinical trials in de novo and recurrence models of GBM are necessary to determine if targeting progenitor GSCs prevents tumour growth and recurrences post treatment. Since mice xenografted with these progenitor GSCs develop tumours faster and exhibit a shorter survival time than mice engrafted with distal GSCs, targeting this most rapidly cycling and functionally aggressive progenitor cancer cell population may be an effective treatment approach.

## Acknowledgements

We would like to thank Carmen Sabau for her contribution to the administrative work of the project, as well as Rosika Bolovan and Maryam Safisamghabadi for technical support. We are grateful to Sara Baig for the design of the schematic in Supplementary Fig. 1a, Greg Chang for supplying reagent for mass cytometry experiments, Jason Karamchandani for help accessing and formatting the TCGA data, Adrian Veres for advice with the initial phase of scRNAseq data processing, and Stefano Stifani for advice with interpretation of the fetal datasets. We also thank the McGill University and Genome Quebec Innovation Centre sequencing platforms, the bioinformatics platform at the Center for Computational Genomics, the Cell Vision Core Facility for Flow Cytometry and Single Cell Analysis, the Imaging Facility, the Histology Facility of the Life Science Complex of McGill University, the Flow Cytometry and Cell-Sorting Facility of the Department of Microbiology and Immunology of McGill University, the Flow Cytometry Facility of the Institut de Recherches Cliniques De Montréal, the Flow Cytometry Facility of the C-BIGR of the Montreal Neurological Institute, the Mass Cytometry Core Facility of the Life Science Complex of McGill University, the Immunophenotyping platform of the RI-MUHC of McGill University. Funding provided by CFI Leaders Opportunity Fund, Genome Canada Science Technology Innovation Centre, Compute Canada Resource Allocation Project and Genome Innovation Node (all to GB and JR), and the Cancer Research Society, Canadian Cancer Research Institute, Brain Tumour Foundation of Canada, and the Canadian Institute of Health Research (all to KP). We wish to thank the TARGiT Foundation, the A Brilliant Night Foundation, and the Argento Family Group Ercole for supporting this work. CPC is supported by the Fonds de Recherche du Québec - Santé Resident Physician Research Career Training Program Phase 1. SA is supported by a George H. Harris Fellowship and a MNI-Fisher Brain Tumour Award. GB is supported by a Fonds de Recherche du Québec - Santé Junior 2 Award. KP is supported by a clinician-scientist salary award from the Fonds de Recherche du Québec - Santé and the William Feindel Chair in Neuro-Oncology. There are no conflicts of interests.

## Author Contributions

C.P.C. and K.P. conceived the project, designed the study, and interpreted the results. K.P. performed the surgeries, and P.L. and X.Y. collected the single cells. D.W. generated the single-cell sequencing libraries. C.P.C. performed scRNAseq-related computational analyses. J.M. performed CNA-related computational analyses. S.A. performed flow and mass cytometry. P.L., S.B., and C.L. performed immunofluorescence imaging. P.L, X.Y., R.A., and S.B. performed the drug and chemotherapy assays. R.A. performed the cloning of the luciferase vector and produced the lentivirus. X.Y., G.R., S.A., and C.P.C. performed the *in vivo* experiments. J.A. and W.Y. provided access to fetal brain samples. M.B., M.C.G., G.B., B.M. and J.R. provided analytical and experimental support. C.P.C., S.A and K.P. wrote the manuscript with feedback from all other authors.

## Methods

### Glioblastoma samples

Glioblastoma samples were harvested under a protocol approved by the Montreal Neurological Hospital’s research ethics board. Consent was given by all patients. Surgeries were performed at the Montreal Neurological Hospital. Pre-operative magnetic resonance imaging was performed for surgical planning. Tumour samples were obtained at the junction of the contrast-enhancing portion of the tumour and brain invasion. In our experience, this location maximizes cell viability, reduces the confounding effects of hypoxia and necrosis, and increases the number of cells which can be extracted from the sample. A certified neuropathologist confirmed all tumour histopathological diagnoses and IDH mutation status by DNA sequencing.

Whole tumour specimens were washed three times in sterile PBS containing penicillin and streptomycin. Specimens were then minced into fragments of less than 1mm in size, before being digested in a collagenase solution containing DNAse and MgCl_2_ for 1-2 hours at 37°C. The digested specimens were washed three times with sterile PBS, and large debris were removed with a 70μm strainer. Residual RBCs were removed using a density gradient in a 1:1 volume ratio with the sample (Lymphoprep, Axis-Shield). Samples were washed five more times in sterile PBS.

### Preparation of the whole tumour and GSC samples

The isolated cells were divided into two parts: one for whole tumour analysis; and one for glioma stem cell enrichment.

Whole tumour cells were prepared for sequencing by first removing endothelial cells and lymphocytes as follows. The isolated cells were resuspended at a concentration of 1e6/mL in PBS. After removing 50μL as unstained control, the live/dead dye, Aqua (Molecular Probes) was added at a concentration of 1:1000. Cells were incubated for 25 minutes on ice, protected from light. Cells were washed once with PBS and resuspended in 100μL of PBS with 1% BSA. FcR block (Miltenyi) was added and incubated for 15min. CD31 conjugated to BV421 (Biolegend), and CD45 (Biolegend) conjugated to PE were added to the suspension at pre-titrated values and mixed well by resuspension and incubated for 25 minutes on ice, protected from light before washing twice with PBS. Compensation beads (Molecular Probes) were used to prepare compensation controls for all antibodies and live/dead used. The sample was then resuspended in PBS with 5% BSA with 25mM HEPES and 2mM EDTA at a final volume of 300-500μL and sorted on the FACS Aria III. Sorted cells were collected in polypropylene tubes with 1mL of icecold FACs buffer with a temperature maintained at 4°C throughout sorting. We selected cells that were negative for CD31 and CD45. Cells were resuspended in PBS with 0.04% BSA for single-cell capture (Supplementary Fig. 1a).

For glioma stem cell (GSC) enriched samples, whole tumour presorted cells were expanded as neurospheres in complete neurocult-proliferation media (Neurocult basal medium containing: Neurocult NS-A proliferation supplement at a concentration of 1/10 dilution, 20ng/ml recombinant EGF, 20ng/ml, recombinant bFGF, and 2μg/ml Heparin) from Stem Cells Technologies. After 7 days of NCC culture, the neurospheres were collected in a tube and spun at 1200rpm for 3 minutes. To dissociate the spheres, Accumax (Millipore) was added to the cell pellet and incubated for 5 minutes at 37°C, they were then washed with PBS, centrifuged and resuspended in PBS with 0.04% BSA for single-cell capture (Supplementary Fig. 1a).

### Human fetal brains

Human fetal brain tissue samples (13–21 gestational weeks) were obtained from the University of Washington Birth Defects Research Laboratory (Seattle, Washington, USA), Centre Hospitalier Universitaire Sainte-Justine (Montreal, Quebec, Canada) and from the University of Calgary (Calgary, Alberta, Canada). These tissues were obtained at legal abortions. The use of the specimens following parental consent was approved by The Conjoint Health Research Ethics Board at the University of Calgary and studies were carried out with guidelines approved by McGill University and the Canadian Institutes for Health Research (CIHR). Cells were freshly isolated as previously described^57^. Briefly, fetal brain tissue was minced and treated with DNase (Roche, Nutley, NL) and trypsin (Invitrogen, Carlsbad, California, USA) before being passed through a nylon mesh. The flow cells were collected in PBS for sorting followed by sequencing (see below).

### Human adult brain

Human autopsy brain specimens were obtained from de-identified excess diagnostic brain tissue that had been slated for incineration. Brains were cut in the coronal plane and immersed in 3% paraformaldehyde (PFA) or formalin for 1–2 weeks and then portions of their lateral ventricular walls were excised. These were further processed for immunolabeling, embedded in paraffin, and 5µm thick sections were cut using a microtome (SLEE).

### Fetal cell sorting

Single-cell fetal cells were washed thrice with excess ice-cold PBS and spun down at 1400rpm for 10min. Cells were resuspended at 1e6/mL of PBS and aqua live/dead dye(Molecular Probes) was added at 1:1000 and incubated for 25 minutes on ice, protected from light. Cells were washed once in excess PBS and were re-suspended at 1e6/40µL and FcR block(Miltenyi) was added at 5µL per 50µL. Cells were mixed well and left to incubate on ice for 15 minutes. CD133-PE (eBioscience), CD45-PerCP/Cy5.5 and CD31-PerCP/Cy5.5 were added at a concentration of 1: 20 and cells were resuspended well before being left to incubate on ice for 25minutes. 1e5 cells were kept aside as unstained control and 5e5 cells were kept aside for fluorescence minus-one gating for CD133 only (FMO-PE).

All cells were washed twice with excess PBS and were spun down at 1400rpm for 5-10 min. Cells were resuspended in ice-cold FACs buffer (5% BSA in PBS with 1% penicillin-streptomycin) before sorting. Sorted cells were collected into polypropylene tubes with 1mL of ice-cold FACs buffer with a temperature maintained at 4°C throughout sorting. All samples were acquired on the BD FACS Aria Fusion III.

Compensation beads (Invitrogen) was used to prepare compensation controls for all antibodies and live/dead stains used. A minimum of 5000 events were acquired for compensation matrix calculation and a minimum of 50e4 total events were collected for fetal samples and analyzed using FlowJo(v10, FlowJo LLC).

### Single-cell RNA sequencing

For each sample, fetal or cancer, an aliquot of cells was taken and stained for viability with calcein-AM and ethidium-homodimer1 (P/N L3224 Thermo Fisher Scientific).

Following the Single Cell 3’ Reagent Kits v2 User Guide (CG0052 10x Genomics) ^29^, a single cell RNA library was generated using the GemCode Single-Cell Instrument (10x Genomics, Pleasanton, CA, USA) and Single Cell 3’ Library & Gel Bead Kit v2 and Chip Kit (P/N 120236 P/N 120237 10x Genomics). The sequencing ready library was purified with SPRIselect, quality controlled for sized distribution and yield (LabChip GX Perkin-Elmer) and quantified using qPCR (KAPA Biosystems Library Quantification Kit for Illumina platforms P/N KK4824). Finally, the sequencing was done using Illumina HiSeq4000 or HiSeq2500 instrument (Illumina) using the following parameter: 26 bp Read1, 8 bp I7 Index, 0 bp I5 Index and 98 bp Read2.

Cell barcodes and UMI (unique molecular identifiers) barcodes were demultiplexed and single-end reads aligned to the reference genome, GRCh38, using the CellRanger pipeline (10X Genomics). The resulting cell-gene matrix contains UMI counts by gene and by cell barcode.

### Analysis of copy number aberrations and isolation of non-cancerous cells

Cells from all samples were pooled in silico. The raw counts of each cell were first normalized using a trimmed mean of M-values (TMM) normalization approach ^58^. This normalization is not affected by outliers but ensures that the majority of the genes support the normalization scale factor. The genome was then tiled by merging consecutive genes into “expressed regions” with a minimum average expression across the cells (5 reads). This new expression matrix was also TMM normalized. For each region and each cell, a Z-score was then computed by subtracting the average expression across cells and dividing by the standard deviation. These Z-scores were winsorized at −3 and 3, minimizing the effect of strong outliers. To focus on the effect of copy number aberrations (CNAs), we minimized expression patterns that are specific to a single expressed region by applying a moving median. Using a sliding window of 7 regions, this moving median approach replaced the expression of a region by the median over the surrounding 7 regions (3 upstream and 3 downstream).

A principal component analysis was performed on the smoothed Z-scores using non-cycling cells (see Cell-cycle and principal components analysis). Because of the genome tiling and moving median, this PCA focuses on expression variability affecting large regions, hence driven by CNA. Louvain clustering was then performed on the K-nearest neighbour graph built using K=100. The similarity between nodes was computed as 1/(1+D) with D the Euclidean distance on the first 20 principal components. ‘KNN’ and a modified version of ‘igraph’ R packages were used respectively for the KNN graph and Louvain clustering ^59, 60^. We scanned the resolution parameters *γ* = 0.1 to *γ* = 1.5, in increments of 0.1. We ran the Louvain clustering 100 times for each resolution, shuffling the order of the nodes in the graph each time. To assess the stability of the clustering at each resolution, we computed the average and standard deviation of the Adjusted Rand Index between pairs of classification 61 (Supplementary Fig. 1c). *γ* = 0.2 was the resolution with the highest average Rand index and lowest standard deviation ^62, 63^. T-distributed stochastic neighbour embedding ^64^, or tSNE, was used to visualize the cells across patients and clusters, using again the first 20 principal components.

Cells were annotated as normal if belonging to one of the two clusters that contained a mixture of cells from different patients (Fig. 1b and Supplementary Fig. 1d). These clusters had low cycling scores and could not be explained by differences in sequencing depth. In addition, these two clusters formed an outgroup when focusing on chromosomes 7 and 10, two chromosomes that are known to host recurrent CNAs in glioblastoma ^19^. As expected, normal cells had lower expression in chromosome 7 and higher expression in chromosome 10. The two clusters of normal cells were remarkable for their expression of lymphocyte genes in one cluster, and oligodendrocyte and endothelial genes in the other (Fig. 1c), which indicates their nature. The presence of two clusters of normal cells is most likely due to subtle cell-type specific patterns that were not fully corrected by the moving median (see example in Supplementary Fig. 7a).

Clones within tumours were defined by running the same Louvain clustering approach separately on the tumour cells of each patient. Here, the number of principal components used were automatically chosen by the ‘quick.elbow’ function of the ‘bigpca’ R package. The optimal resolution gamma was chosen as described above (Supplementary Fig. 7b). When the best average Adjusted Rand Index was lower than 0.7, we considered the clustering too unstable and grouped all the cells from the patient into one unique clone. To characterize the CNA profile of each clone, cells were merged into super-cells by summing their raw gene counts. For each clone, we created 10 super-cells, each by merging 30 randomly selected cells. Super-cells from normal cells were created similarly and were used later as baseline. The super-cells for each clone were then pooled, TMM normalized genes were merged as above to create expressed regions with at least 20 reads on average. For each expressed region and each super-cell, a log-ratio was computed by dividing the normalized counts by the average counts in the normal super-cells. Using the log-ratios and a multivariate Gaussian mixture hidden Markov model (HMM), regions were classified as loss, neutral, or gain. The HMM had three states with means log(0.5), 0 and log(1.5), the empirical standard deviation estimated from the data, and represented the 10 super-cells simultaneously for each clone. The ‘viterbi’ function from ‘RcppHMM’ R package was used to estimate the most likely states of a “GHMM” object. The transition probability was set to 10^−40^. We define a loss (gain) of a chromosome if more than 50% of the regions are in the loss (gain) state. Finally, the significance of each chromosomal CNA was confirmed using a Wilcoxon test on the median chromosome expressions. All the CNAs showed p-values below 0.001. The HMM analysis was also run on the normal cells and no CNAs were detected. We used a Chi-squared test to compare the cell distribution across the 4 TCGA signatures between pairs of clones (Supplementary Fig. 2a). Except for the first clone of BT333, clones had significantly different TCGA signatures (p<0.01).

### Signal processing for transcriptional data

Low complexity cells (< 2000 genes or <1800 UMI detected), dying cells (> 12% UMI to mitochondrial genes, Supplementary Fig. 8a), non-cancerous cells (see Analysis of copy number aberrations and isolation of non-cancerous cells) and genes with no counts were removed from the analysis. Next counts were adjusted in each cell according to a size factor akin to TPM. Genes which accounted for more than 1% of UMI in a given cell were not counted towards the UMI sum of this cell. Similar to previous studies ^27, 28^, each cell was normalized to 1e5 UMI.

Signal-containing, non-random genes were selected in each sample. This was done in a manner to that described by Klein *et al.* ^27^. Briefly, we selected genes with a variance that was most likely to exceed that of a Poisson using a Fano statistic. We calculated the Fano factor for each gene in the matrix M. We selected genes above a selected threshold of Fano statistic. The threshold was selected to obtain around 3000 highly variable genes in every sample. We then applied a base 2 logarithm to obtain the normalized expression matrix. A z-score by gene was applied at this point for single sample analyses. For analyzes spanning multiple samples, we combined the normalized expression matrices on the basis of the intersection of their significant genes. Z-score across all cells and samples was applied by gene thereafter.

### Filtering the fetal brain and cancer samples

We removed ependymal cells and microglia from later fetal analyses. These were seen as separate clusters in PC1 and PC2 in most samples. Microglia had high expression of genes such as P2RY12 and CX3CR1^65–67^, while ependymal cells had high expression of SPAG6, FOLR1, and FOXJ1 ^68–70^.

BT346 contained many cells with a signature not seen in other samples. These clustered separately in tSNE and PCA. We used k-means (k=2) to separate them from the other cells. A gene set enrichment analysis (GSEA, see Quantification and Statistical Analysis for methodology) showed that the top 4 most significant gene sets were linked to hypoxia (e.g. “HALLMARK_HYPOXIA”, “MENSE_HYPOXIA_UP”). This tumour was unique in that the MRI region of contrast enhancement was very thin. It is thus likely that some cells from the necrotic core were isolated. We excluded the hypoxic cells in BT346 from later analyses and did not include them in the total cancer cell number reported.

### Cell-cycle and principal components analysis

We positioned all cells within the cell-cycle according to the method presented by Tirosch *et al.*^31^. Briefly, each cell obtained a score for the G1/S phases and a score for the G2/M phases (Supplementary Fig. 8b). A list of genes deemed characteristic of those cell-cycle states was used^31^. Each score was defined as the sum of the expression of all genes within its corresponding gene set, then z-scored across cells. Since most cells are not cycling (Supplementary Fig. 8b), we defined non-cycling cells as those with both G1/S and G2/M scores less than 0.

Cycling-free principal components analysis (PCA) was performed for each sample individually as follows. The PCs, or eigenvectors of the covariance matrix, were obtained from the non-cycling cells only (as defined above). We then use these cell-cycle-independent eigenvectors to project the complete dataset in PCA space (Supplementary Fig. 1e).

The first PC of every GSC sample was highly conserved (see Results). To quantify this, we compared the ranking of genes by PC1 loadings across samples. The actual ranking of each gene was obtained in all samples. To obtain the expected ranking, the actual rankings were averaged by gene, and these averaged values were then ranked. For each gene, we thus obtained five actual rankings (one per sample) and one expected ranking. R^2^ was obtained by least-square linear regression in PC1 and PC2 separately (Matlab, *fitlm*).

### Classifying cells by TCGA subtype

TCGA subtype for each whole tumour cell was obtained by scoring each cell for their proximity to each TCGA centroid^19^. The highest score obtained by a given cell defined the subtype of the cell. We used this method on the original TCGA dataset and found we could correctly classify 89.7% of all tumours (data not shown).

Proximity is calculated as follows. The position of a cell in the TCGA transcriptomic space (*X_ij_*) is obtained from the expression of the genes present in the TCGA signature (*S*). The unit vector of this cell’s position is then projected onto the unit vector of the signature of interest using a dot product.

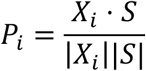

where *P* is the projection score and *S* is the signature of interest.

### Community detection in fetal samples

To properly cluster fetal cells in cell types, the modular structure of the gene coexpression network was estimated using community detection. Data from all fetal samples were merged as explained above. PCA was performed on the merged dataset (see Principal components analysis above). The first ten PCs were selected based on the importance of their corresponding eigenvalue (Supplementary Fig. 8c). The connection weights were computed as 1/(1+D) with D as the Euclidean KNN graph between nodes in this PC space, with K=50. Self-weights were set to 0 to promote the formation of communities.

Again, the goal of the analysis was to identify groups of cells that are more similar to each other than other cells. This constraint was operationalized in terms of the modularity *Q* ^59^:

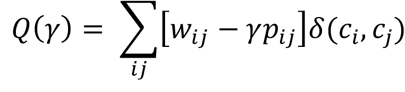

where *w_ij_* is observed connection weight between nodes *i* and *j*, while *p_ij_* is the expected connection weight between those nodes. The Kronecker delta function, *δ(c_i_, c_j_)* is equal to 1 when nodes *i* and *j* and assigned to the same community (*c_i_ = c_j_*) and zero otherwise (*c_i_ ≠ c_j_*), ensuring that modularity is only computed for pairs of nodes belonging to the same community. The resolution parameter *γ* scales the importance of null model *p_ij_*, potentiating the discovery of larger (*γ* < 1) or smaller communities (*γ* > 1)^60^.

In the present study, the expected connection weight between pairs of nodes was defined according to a standard configuration model, such that:

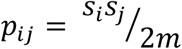

where *s_i_* = Σ*_i_ w_i_* is the strength of node *i* and *m* = Σ*_i, j>1_ w_ij_* is total weight of connections. Under this null model, communities are considered to be of high quality if the constituent nodes are more highly correlated with each other than in a randomly rewired network with the same strength distribution and density.

The quality function *Q* was maximized using a Louvain-like locally greedy algorithm ^71^, as implemented in the Brain Connectivity Toolbox (*community_louvain.m*) ^72^. We scanned the resolution parameters *γ* = 0 to *γ* = 1.5, in increments of 0.1. At each scale, the Louvain algorithm was run 100 times to find a partition that maximized the modularity function ^71^.

To select an appropriate scale, we computed the z-score of the Rand index between all pairs of partitions at each scale^61^. We selected the resolution at which the mean pairwise Rand index to standard deviation ratio (signal-to-noise ratio - SNR) was greatest across the partition ensemble^62, 63^. The logic behind this approach is that if there exists a particularly well-defined community structure at some topological scale, then it should be relatively easy to detect, and the partitions will not vary greatly across runs. Supplementary Fig. 3b shows the SNR of all pairwise Rand indices. Based on this method, we selected *γ* = 1.0. Once the scale was selected, we used the consensus heuristic described by Bassett et al. ^62^ to find the most representative partition in the ensemble (Brain Connectivity Toolbox; *consensus_und.m*).

We further studied cluster 2, which expressed glial markers of both astrocytic and oligodendrocytic origin. The algorithm described above was used again, with resolution parameters scanned from *γ* = 0 to *γ* = 1.5, in increments of 0.01. Lower increments were used because less total nodes allowed for additional computational time. *γ* = 0.44 was the consensus or most representative partition.

The values shown in the similarity matrix heatmap (Fig. 3b) are the inverse of the diffusion pseudotime^38^ between cells.

### Differential expression of fetal cell types

We assessed differentially expressed genes between fetal brain cell types by comparing each cell type to all other combined. A Mann-Whitney U test (Matlab, *ranksum.m*) was applied on the log expression value (before z-score) of each gene sequentially. P-values were adjusted for multiple testing using approach described by Storey ^73^, and reported as q-value (Matlab, *mafdr.m*).

### Creation of the fetal roadmap

We aimed to create a fetal roadmap, or a transformation of transcriptomic space descriptive of the transitions that exist between the cell types present in glioblastoma. Simply, we first determined which fetal cell types were most representative of the cancer; then we created a PC space of these cell types onto which the cancer could be mapped.

We determined the most representative cell types by finding each cancer cells closest transcriptomic fetal brain cell neighbour. Data from all whole tumour and fetal brain samples were merged as described above. Each fetal cell type was randomly subsampled (Matlab, *randsample.m*) to obtain an equal number of cells for each cell type. Whole tumour samples were similarly subsampled. The top ten PCs for this new fetal dataset were calculated (see Removal of Cell Cycle section), and both fetal brain and cancer datasets were projected in this space. The closest fetal brain cell neighbour for each cancer cell was found (Matlab, *knnsearch.m*). We refer to this as a capture of this cancer cell by the fetal cell. The number of cancer cells captured by each fetal cell type was tabulated. Neuronal progenitor cell types were tabulated under their more differentiated counterpart.

Four fetal brain cell types were retained for the creation of the cancer roadmap. As was done above, fetal brain cells from these four subtypes were randomly subsampled to balance their numbers. Genes common to both cancer cells and fetal brain cells were kept for the analysis (n=398 for whole tumour, n=401 for GSC, and n=345 for GSC and whole tumour combined). Cell-cycle-free PCA was performed on these fetal brain cells (see above) and four PCs were kept. These four PCs were sufficient to appropriately resolve all four fetal brain cell types included. We defined this as the roadmap. To refine this separation and better capture the transitional nature of this data, we performed diffusion embedding on the roadmap. Briefly, from the roadmap we calculated a transition matrix (REF, *diffusionmap.T_nn.m, k* = 50, *nsig =* 10). The top four eigenvectors were obtained and normalized (ref, *eig_decompose_normalized.m*). The first eigenvector was dropped as the steady state of the transition matrix. Eigenvectors 2 to 4 were defined as DC1, DC2, and DC3, respectively. The glial progenitor score was defined as DC3.

### Mapping of cancer cells to the fetal roadmap

The aim of the roadmap was to highlight the underlying hierarchical organization while de-emphasizing interpatient variability. Hence, we projected cancer cells (whole tumour, GSCs, or both) onto the 4-dimensional fetal PC space of the roadmap. This represents the mapping of cancer cells onto fetal PC space. We used these results to obtain the diffusion and the simplified PC mapping of cancer, as we will explain below.

To obtain cancer mapping in diffusion space, we first obtained the transition matrix of the fetal roadmap as described above. From the PC cancer mapping, a separate transition matrix was obtained for cancer but solely as a function of the fetal brain cells (*diffusionmap.T_nn.m, k* = 50, *nsig* = 10). This amounts to obtaining the transition matrix of the combined fetal brain/cancer data but only one cancer cell at a time. The cancer transition matrix (*T_cancer_*) was then projected onto the roadmap diffusion components (*ϕ_fetai_*) defined above.

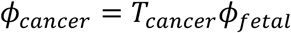

The resulting diffusion component vectors (*ϕ_cancer_*) represented the mapping of cancer cells in the roadmap diffusion space.

To rule out the possibility that this hierarchical distribution could be the product of chance, we created control cells by randomly swapping the genes in our whole tumour cells. These control cells would have had the same depth of sequencing, but gene signatures were absent. Using a Kolmogorov-Smirnov statistic, we found that our cancer cells and control cells had a very significantly different distribution when projected on the roadmap, both in diffusion space and PC space (p-value < 1e-22 for both).

Next, we sought to create a simplified PC roadmap, in an effort to better capture biological relevance. This is because both GPC and OLC populations contain progenitors, and a reliable surface marker to differentiate the two was not found. In the PC roadmap, PC2 and PC3 separate fetal OPC and fetal GPC from the other cell types, respectively. Therefore, to make a combined progenitor score, we summed the values of PC2 and PC3 (Fig. 5c). PC1 already separated astrocytes (positive values) from interneurons (negative values). We defined the latter as our lineage score. Cancer genes which correlated with each of these two scores (see Table 3) guided our search for markers for each cell type.

### Classification of cancer cells

In order to compare the signatures of cells at the extreme ends of the hierarchy, we aimed to classify the cancer cells by cell type. Using the annotated fetal data in diffusion roadmap space as a training set, we performed a linear discriminant analysis (LDA, Matlab, *fitcdiscr.m*). Cancer cells in diffusion roadmap space were classified using this model (Fig. 4b). In order to classify extremes of the hierarchy only, any cell with a probability of incorrect classification of more than 0.01% was left unclassified.

### Pathway enrichment for progenitors in whole tumour

Whole tumour cells classifications were obtained using the LDA method described above. Progenitor and astrocytic classifications were used. As had been done previously ^41^, each gene was ordered according to its signal to noise ratio (SNR) for the progenitor vs the astrocytic cell type

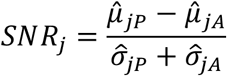

where 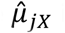 is the estimated mean log expression of gene *j* for progenitor (*P*) and astrocytic (*A*) cancer cells; and 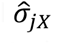 is the estimated standard deviation of log expression for gene *j*. A Mann-Whitney U test (Python, *scipy.stats.mannwhitneyu*) was used to determine if the *SNR* values for genes in a given gene set were significantly different than the *SNR* not in this gene set (see Supplementary Fig. 8d for an example). All gene sets in the *c2.all.v6.0* dataset from the Broad Institute ^41–43^ were tested, using the genes present in our combined whole tumour dataset (n=970, see Table 2 for list of genes). P-values were adjusted for multiple testing using the approach described by Storey ^73^, and reported as q-value.

### Mass cytometry

Metal tagged mass cytometry antibodies were purchased from Fluidigm. Where tagged antibodies were not available, purified antibodies lacking carrier proteins were labelled with heavy metal loaded maleimide conjugated DN3 MAXPAR chelating polymers (Fluidigm) according to the recommendations provided by Fluidigm.

Cells were stained according to a well-established protocol for cell cycle staining ^39^. Briefly, cells were incubated with IdU at 50µM final concentration for 30min at 37°C and 5% CO_2_ in stem cell media. A live/dead stain was performed by incubating cells with 5µM cisplatin (Fluidigm) at room temperature for 5 minutes. Cells were washed twice with cell staining buffer (CSB), composed of standard phosphate-buffered saline (PBS) with 0.5% BSA and 0.02% sodium azide, twice. Before cell surface antibody labeling, Fc-receptors were blocked using human BD Fc block (BD biosciences). Cells were then labeled with a surface antibody panel which included CD9, CD24, CD44, CD133, PDGFRa, HLA-ABC, Olig2 and CD45 and CD31 and incubated on ice for 25min. Cells were then washed and fixed using Fix I buffer (Fluidigm) for 15min. This was followed by two more washes with CSB and ice-cold methanol fixation for 15min on ice. Intracellular labeling was carried out for 25min on ice. A final two more washes with CSB were carried out followed by an overnight incubation in Fix and Perm buffer (Fluidigm) with 125nM of iridium intercalating dye (Fluidigm).

Mass cytometry data were analysed using FlowJo (v.10, FlowJo LLC) and a hyperbolic arcsine transformation on all parameters.

### Glioma stem cell sorting

Multiparametric flow cytometry was carried out by labeling cells with CD9 preconjugated with BV421 (BD Pharmingen), CD24 preconjugated with APC or APC-H7 (Miltenyi), CD44 preconjugated with AF700 (BD Pharmingen), and CD133/PROM1 preconjugated with PE or PE/Vio770 (eBioscience and Miltenyi). After leaving aside 1e5 cells as unstained control, cells were resuspended in PBS at a concentration of 1e6/mL. Aqua live/dead dye (Molecular Probes) was added at 1:1000 and incubated for 25 minutes on ice, protected from light. Cells were washed and 1e5 cells were kept aside for fluorescence minus-one (FMO) controls and 1e6 cells were used for complete staining with antibodies. FMO controls were prepared for all colours except aqua (live/dead). All cells were completely stained with antibodies at a final dilution of 1:50-1:20. FMO controls were used to identify for positive/negative staining.

Sample preparation post-staining for sorting and data acquisition was carried out as described above.

### Luciferase vector

The Red Firefly Luciferase sequence was amplified from the pCMV-RedFLuc (Targeting Systems, CA, USA) and cloned into the bidirectional EF1/PGK promoter lentiviral vector (System Biosciences, Palo Alto, USA). The final construct was named PGK-GFP-LUC. Lentivirus was produced as per the protocol described by Ritter *et al*.^74^. Expression of the construct was validated by luciferase assay and fluorescence microscope.

### Mouse xenotransplantation

All animal procedures were approved by the Institution’s Animal Care Committee and performed according to the guidelines of the Canadian Council of Animal Care. We orthotopically injected 100k (for general tumourigenicity and E2F inhibition) or 5k (cluster tumourigenicity) GFP^+^/Luciferin^+^ GSCs into female NOD-SCID gamma mice (Charles River, Wilmington, USA), as previously described ^75, 76^. Briefly, mice were anesthetized at 5 weeks of age using isofluorane (Fresenius Kabi, Bad Homburg, Germany) and placed on a stereotaxic apparatus. A midline scalp incision was made and a burr-hole (3mm) was created 2.2mm lateral to the bregma using a high-powered drill. The injection needle of an Hamilton syringe (Hamilton, Reno, USA) was then lowered into the burr-hole to a depth of 2.5mm and cells were transplanted into the striatum. Animals were frequently monitored and then euthanized at the appearance of distress signs and/or 10% weight decrease. These animals were perfused with phosphate-buffered saline and their brain collected. Kaplan-Meier curves were created according to the survival results. A Cox proportional hazard ratio model was used to assess significance, with patient cell line and cell type (Fig. 6e) or treatment group (Fig. 7f) as covariates. This analysis was performed in R using the packages *splines* and *survival*.

Harvested brains were placed in 10% neutral buffered formalin for 72 hours at room temperature. After formalin fixation, specimens were processed and paraffin-embedded. Five μm tissue sections were prepared and mounted on a poly-L-lysine-coated glass slides for subsequent analysis.

### In vivo imaging

To monitor tumour growth, we imaged each mouse every 2 weeks using the In Vivo Imaging System (IVIS) Sprectrum (Perkin Elmer, Waltham, USA) according to the manufacturer’s instructions. Briefly, we intraperitoneally injected a solution (15mg/ml) of luciferin (Perkin Elmer) at the dose of 150mg/kg, and after 3 min, mice were anesthetized using isofluorane. At 10 minutes from luciferin injection, we positioned the mouse in the imaging system and began image acquisition. The exposure time was automatically determined by Living Image 4.5.2 software (Perkin Elmer). Results are reported as number of photons emitted, and a 2-sample student-t test was performed.

### Immunofluorescence

GSCs were grown on laminin (10µg/ml) coated coverslips, and fetal neural stem cells were grown on Matrigel in the supplemented mTeSR1 basal medium (STEMCELL Technologies). Both were fixed with 3% PFA and permeabilized with 0.5% TritonX-100 before being immunolabeled with indicated antibodies followed by secondary antibodies. Cover slips were mounted on glass slides using ProLong™ Diamond Antifade Mountant with DAPI (Invitrogen) to counterstain cell nuclei. Fluorescent images were acquired using ZEISS LSM 700 laser scanning confocal microscope with a 20X or 63X objective.

For the GSC assays, the total number of Ki67^+^ cells relative to total cell number were quantified from 10 fields for each patient cell line (n=3).

For tissue sections (brain and tumour) immunohistochemistry, samples were baked overnight in a standard laboratory oven at 60 degrees, then deparaffinised and rehydrated using a graded series of xylene and ethanol, respectively. Antigen retrieval was done using citrate buffer (pH 6.0) for 10 or 20 minutes at 120 °C in a decloaking chamber (Biocare Medical). The slides were then blocked for 20 minutes with a commercial protein block (Spring Bioscience), incubated overnight at 4°C with indicated antibodies, then slides were washed with IF buffer (PBS + 0.05% tween20 + 0.2% triton X-100), following by incubation (1h at room temperature) with according secondary antibodies (Invitrogen). Cover slips were mounted on glass slides using ProLong™ Diamond Antifade Mountant with DAPI (Invitrogen) to counterstain cell nuclei. Fluorescent images were acquired using ZEISS LSM 700 laser scanning confocal microscope with a 63x objective.

For tumours, total number of CD133^+^ or Ki67^+^ or both CD133^+^ and Ki67^+^ cells relative to total cell number were quantified from at least 10 images from each patient. A χ^2^ test was performed to obtain the level of significance. A significant association of Ki67 and CD133 was found in all patients.

Primary antibodies used: anti-GFAP (Abcam); anti-Olig2 (EMD Millipore), anti-Ki-67 (Invitrogen and Abcam), and anti-CD133 (Miltenyi Biotec).

### Chemotherapy and targeted therapy assays

Temozolomide (TMZ) – GSCs from each cluster type were seeded on laminin (10µg/mL, Sigma) at a concentration of 10 000 cells/well in a 96-well plate and were subsequently treated for 5 days with varying concentrations of TMZ (Sigma Aldrich) ranging from 1µM to 750µM. 50μL of XTT was prepared according to the manufacturer’s instructions (Life Technologies), and the XTT mix solution was added to the cells and further incubated for 3 hours at 37°C. The absorbance at 450nm was measured on an Epoch Microplate Spectrophotometer (Biotek Instruments, USA).

HLM006474 **–** GSCs from each cluster type were plated on laminin (10µg/mL, Sigma) at 5000 cells/well in 96 well overnight in culture media. The following day, HLM006474 (or DMSO) was added to a 10µM final concentration in a final volume of 200μL. Following 7 days of incubation at 37°C, an XTT was performed as described above.

Combination therapies – GSCs were plated on laminin (10µg/mL, Sigma) at a concentration of 10 000 cells/well in a 96-well plate and treated with either TMZ (50µM) for 6 days, HLM (7µM) for 6 days, or HLM006474 (7µM) for 3 days followed by TMZ (50µM) for 3 days. After these 6 days of treatment at 37°C, an XTT assay was performed as described above. Sphere forming assay – GSCs from each cluster type were plated at 150 000 cells/well in 6 well plates with 20µM HLM006474 in a final volume of 3ml. After 7 days, cells were imaged with 10x objective with Invitrogen EVOS FL/FL color microscope. Sphere diameter measurements were made with Image J. 6502 spheres were measured in two different patient GSC cell lines. An arbitrary cut off for big and small spheres was set at 65µm. A multivariate logistic regression was used to assess the likelihood of finding big spheres in each of the different GSC cell types treated, using patient cell line and cell type as variables. There was no significant difference between patient cell line (p=0.69). Error bars reported in Fig. 7c are the 95% confidence interval associated with each OR.

All assays were performed in 3 different patient cell lines in 3 or more different cell passages and 5 technical replicates. P-values describe differences in cell types and were calculated using a two-sample t-test. Stock solutions of TMZ (Sigma-Aldrich), HLM006474 (Tocris-Bioscience) were prepared in dimethyl sulfoxide (DMSO; Sigma-Aldrich), and were added to cells for a final DMSO concentration of <0.1%.

### Code availability

All computations and quantifications were performed using Matlab, R, and Python programming languages and are available upon request.

